# Co-occurrence of ST412 *Klebsiella pneumoniae* isolates with hypermucoviscous and no-mucoviscous phenotypes in a short-term hospitalized patient

**DOI:** 10.1101/2023.06.20.545774

**Authors:** Qinghua Liang, Biying Zhang, Wei Wang, Nan Chen, Jingjing Luo, Ying Zhong, Feiyang Zhang, Zhikun Zhang, Alberto J. Martín–Rodríguez, Ying Wang, Li Xiang, Jiaru Zhuang, Renjing Hu, Yingshun Zhou

**Affiliations:** Department of Pathogen Biology, School of Basic Medicine, Public Center of Experimental Technology of Pathogen Biology Technology Platform, Southwest Medical University, Luzhou, 646000, China; Department of Clinical Sciences, University of Las Palmas de Gran Canaria, 35016, Las Palmas de Gran Canaria, Spain; Department of Microbiology, Tumor and Cell Biology, Karolinska Institutet, 17165, Stockholm, Sweden; Wuxi School of Medicine, Jiangnan University, Wuxi, Jiangsu, China; Department of Laboratory Medicine, Jiangnan University Medical Center, Wuxi, China

**Keywords:** *Klebsiella pneumoniae*, whole genome sequencing(WGS), RNA-Seq, hypermucoviscosity, non-synonymous mutations

## Abstract

Hypermucoviscosity(HMV) is a phenotype that is commonly associated with hypervirulence in *Klebsiella pneumoniae*. The factors that contribute to the emergence of HMV subpopulations remain unclear. In this study, eight *K. pneumoniae* strains were recovered from an inpatient who were hospitalized for 20 days. Three of the isolates exhibited a non-HMV phenotype, which was concomitant with increased biofilm formation and higher siderophore secretion than the other five HMV isolates. All eight isolates were highly susceptible to serum killing, albeit HMV strains were remarkably more infective than non-HMV counterparts in a mouse model of infection. Whole genome sequencing(WGS) showed that the eight isolates belonged to the K57-ST412 lineage. Average nucleotide identity(ANI) analysis indicated that eight isolates share 99.96% to 99.99% similarity and were confirmed to be the same clone. Through comparative genomics analysis, 12 non-synonymous mutations were found among these isolates, seven of which in the non-HMV variants, including *rmpA*(R96G) and *wbap* (S435R), which are assumed to be associated with the non-HMV phenotype. The mutations *manB*(G440L), *dmsB*(R193W) and *tkt*(A643N) occurred in HMV isolates only. RNA-Seq and RT-qPCR revealed transcripts of genes involved in transporter activity, carbohydrate metabolism and energy metabolism, including *cysK*, *paaF*, *vasD*, *celC* and *fruA*, to be significantly dysregulated in the non-HMV strain K201060 compared to the HMV strain K201059, suggesting a participation in HMV phenotype development. This study suggests that co-occurrence of HMV and non-HMV phenotypes in the same clonal population may be mediated by mutational mechanisms as well as by certain genes involved in transport and central metabolism.

**Importance:** *K. pneumoniae* with a hypermucoviscosity(HMV) phenotype is a community-acquired pathogen that associated with increased invasiveness and pathogenicity, and underlying diseases are the most common comorbid risk factors inducing metastatic complications. HMV was earlier attributed to the overproduction of capsular polysaccharide, and more data point to the possibility of several causes contribute to this bacterial phenotype. Here, we describe a unique event in which the same clonal population showed both HMV and non-HMV characteristics. Studies have demonstrated that this process is influenced by mutational processes and genes related to transport and central metabolism. These finding provide fresh insight into the mechanisms between behind co-occurrence of HMV and non-HMV phenotypes in monoclonal populations as well as potentially being critical in developing strategies to control the further spread of HMV *K. pneumoniae*.

## Introduction

*K. pneumoniae* exhibiting phenotypic hypermucoviscosity(HMV) is frequently associated with hypervirulence. The emergence of HMV, hypervirulent and multidrug resistant isolates is rising global concerns and a gradual increase in the morbidity and mortality of K. pneumoniae infections(1), which include urinary tract infections, meningitis, pyogenic liver abscesses, and empyema, among others(2).

HMV is a prominent phenotypic feature characterized by the formation of a viscous filament >5mm when the colony is stretched by the inoculation loop on an agar plate(3). The correlations between the HMV phenotype and certain specific regulatory factors, is widely accepted. For instance, the mucoid regulator gene, *rmpA,* located on a large virulence plasmid, has been shown to be required for HMV phenotype development and as such, deletion of *rmpA* resulted in reduced amount of capsule and a non-HMV phenotype(4). HMV has also been attributed to capsular polysaccharide overproduction, albeit it does not fully account for the presence of a HMV phenotype(5). The majority genes in the capsular polysaccharide(*cps*) cluster, including *wza, wzc, wzy,* and *wckU,* are involved in the transport and assembly of capsules polysaccharide(6). A non-HMV phenotype due to mutations in the *cps* cluster has been shown to occur among clinical carbapenem-resistant *K. pneumoniae* isolates, affecting susceptibility to carbapenems but also resulting in a better ability to form biofilm (7).

HMV and CPS overproduction are energy intensive processes(8), implying that a number of genes associated with the TCA cycle and cellular energy metabolism, might mediate the HMV phenotype. Recent analysis of a library of transposon integration mutants within the hypervirulent strain KPPR1 suggested that a number of disrupted metabolic genes have diverse effects on HMV and CPS production(9). Thus, six genes induced HMV loss and CPS reduction(*galU, rfaH, wzyE, arnD, arnE* and *wcaJ*), five resulted in HMV loss but did not affect CPS content(*uvrY, miaA, galF, arnF* and *orf2*) and three reduced CPS content but did not affect HMV(*rnfC2, arcB* and *pgm*)(9). These data suggest that metabolism-related genes are associated with the synthesis of HMV. Therefore, mutations or alterations in the expression of these genes with similar functions might contribute to the emergence of HMV. In addition to metabolism, other factors influence the HMV phenotype, such as antibiotic stress or the host immune system, suggesting that the factors influencing HMV are diverse(10). The full breadth of determinants behind HMV formation is yet unknown and their investigation is essential for an improved prevention and control of *K. pneumoniae* infections.

In this study, eight ST412-K57 *K. pneumoniae* isolates derived from the same clonal population were obtained from a short-term hospitalized patient and the HMV phenotypes were investigated. Whole genome sequencing(WGS), RNA-Seq, CPS extraction, serum killing assay, biofilm formation, siderophores secretion and virulence tests were performed to elucidate the factors contributing to HMV evolution of *K. pneumoniae in vivo* in patients, as well as the selective advantage of HMV phenotypic variation in terms of biofilm production and siderophores biosynthesis.

## Results

### Case presentation and bacterial isolates

On 25 April 2020, a 75-year-old woman was admitted to the Affiliated No.2 People’s Hospital in Wuxi, China, for convulsion and unconsciousness. The previous medical records indicated that the patient has underlying conditions (diabetes mellitus and hypertension). Laboratory data revealed a WBC count of 24000/mm^3^, C-reactive protein level of 329mg/L. A presumed diagnosis of septic shock was made. The inpatient was treated with the empirical administration of cefoperazone sodium and sulbactam sodium. On the next day, two *K*. *pneumoniae* stains (K201054 and K201055) were isolated from blood culture and one *K*. *pneumoniae* strain(K201047) was isolated from cerebrospinal fluid(CSF). Meropenem was administered for the treatment of the infections. During hospitalization, an additional five K. pneumoniae strains were isolated, two from feces, two from urine, and one from a CSF specimen(Fig.1). After symptomatic treatment, the inpatient was finally cured and discharged. Antimicrobial susceptibility testing showed that the eight *K*. *pneumoniae* isolates were susceptible to the tested antibiotics tested (ceftazidime, cefotaxime sodium, ciprofloxacin, chloramphenicol, tetracycline, meropenem and polymyxin).

**FIG 1.**
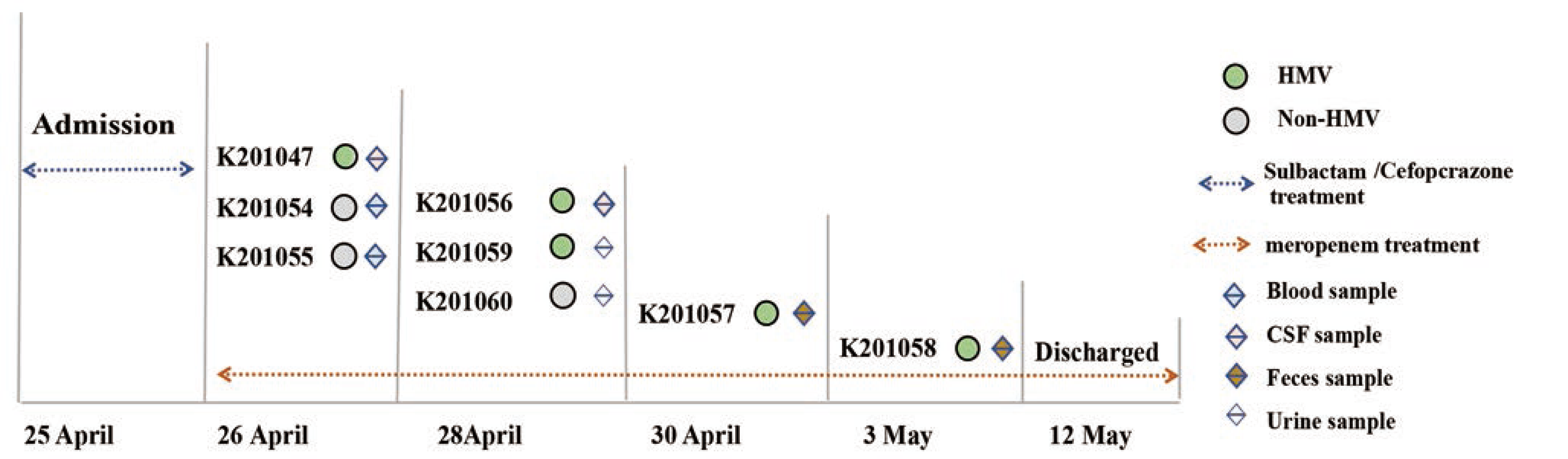
Timeline of the eight episodes of *K*. *pneumoniae* infections in the inpatient. The diamonds of various colors represent the specimen source. Grey circles represent the non-HMV phenotype, and green circles indicate the HMV phenotype. The dotted line represents the antibiotic used for treatment.

### CPS content is associated with the HMV phenotype

To investigate the HMV phenotype of the eight *K. pneumoniae* isolates retrieved during the infection episode, we performed the string test on colonies grown on agar plates. This test revealed that five isolates were HMV (K201047, K201056, K201057, K201058, K201059), whereas the other three were non-HMV (K201054, K201055 and K201060) (Fig.2A). HMV is associated to poor sedimentation. Therefore, to validate the string test observations, we performed a natural sedimentation assay in which bacteria were allowed to sediment by gravity. The supernatant of HMV strains remained turbid with a median OD_600_-supernatant/OD_600_-total ratio of approximately 0.5, whereas that of non-HMV isolates did not exceed 0.2(Fig.2B). In addition, given that the majority of HMV strains have the ability to overproduce capsule, we next extracted the capsular polysaccharides. In agreement, the capsule content of the HMV isolates was significantly higher compared to that of non-HMV isolates (*p*< 0.05) (Fig.2C). Taken together, our results confirmed K201047, K201056 and K201057, K201058, and K201059 to be HMV isolates exhibiting copious CPS production.

**FIG 2.**
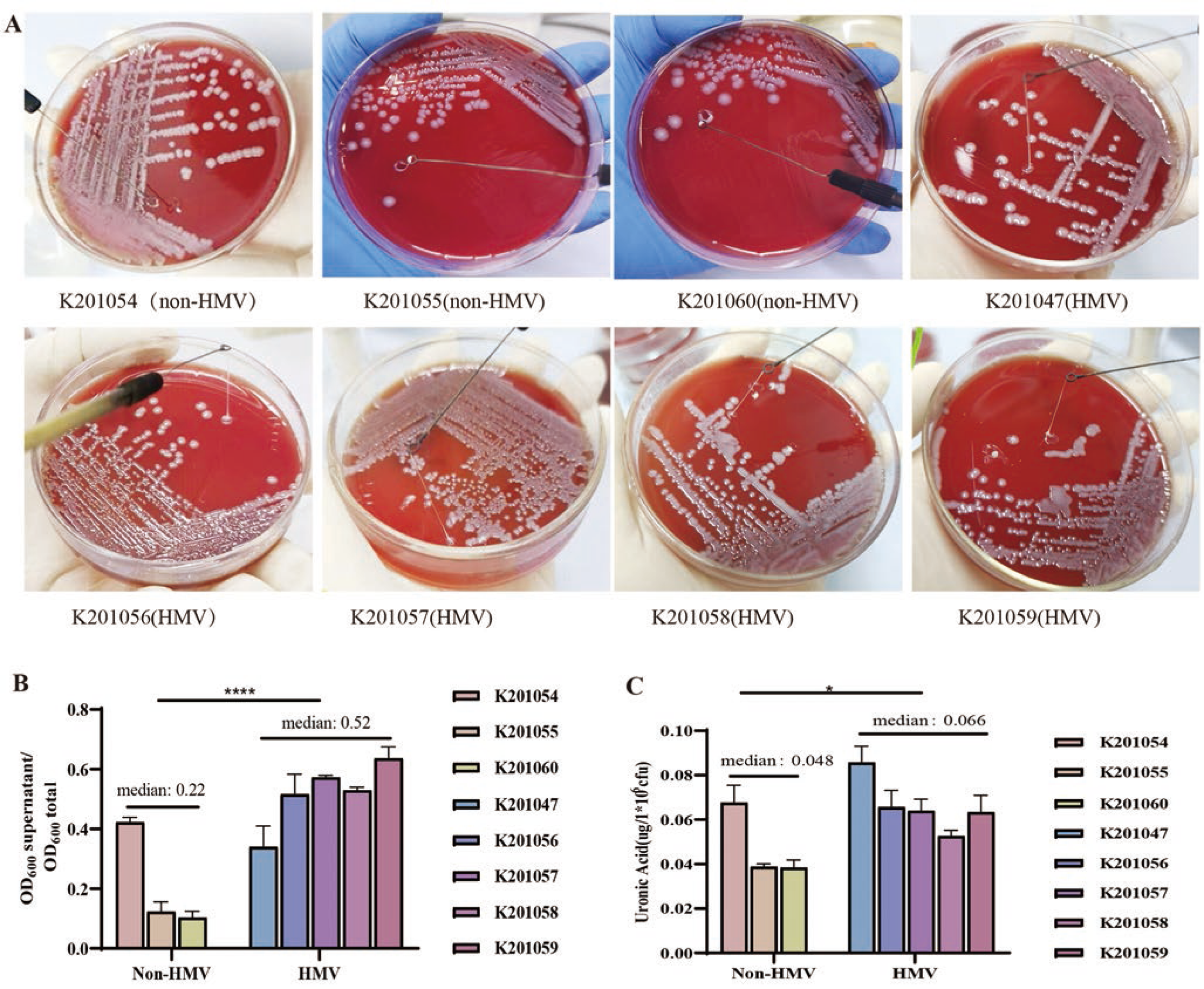
The mucoviscosity phenotype and capsule production of the eight clinical isolates. **(A)**Colony morphology and string test. **(B)** Sedimentation properties of the isolates. **(C)** Amounts of CPS produced as determined by the uronic acid assay. Each assay was independently repeated at least three times. The data represent the mean ± SD and asterisks indicate significant differences (* p < 0.05, **** p < 0.0001) between the compared groups using *t-test*.

### Non-HMV strains show reduced pathogenicity and increased biofilm formation

To investigate the association between phenotypic HMV and other properties relevant to *K. pneumoniae* infectivity and persistence, we first evaluated whether the different mucoid phenotypes affected the ability of the isolates to biofilm formation. It was found that the non-HMV clinical isolates had higher biofilm-forming capacity than HMV isolates, a 3-fold reduction in total biofilm biomass with respect to the former as determined by crystal violet staining(Fig.3A). Production of siderophores such as yersiniabactin, aerobactin, or salmochelin, are known to contribute to *K. pneuominiae* virulence(11). To investigate differences in siderophore production between HMV and non-HMV strains, we performed a chrome azurol S (CAS) agar assay, in which siderophore production is phenotypically determined by the generation of an orange halo. In this test, non-HMV isolates had a greater capacity for siderophore secretion than HMV isolates (p<0.01), as assessed by the area of their halos (Fig. 3B). Next, we tested whether the HMV phenotype offered protection against serum-mediated killing. As shown in Fig.3C, all isolates were completely eradicated after three hours of co-incubation with human serum, clearly indicating that all isolates were highly susceptible to serum killing. To evaluate the pathogenicity of HMV isolates and non-HMV isolates *in vivo*, a mouse model of intraperitoneal infection was employed, In the liver, spleen and kidneys of infected animals, the non-HMV isolates had colonization levels about 3 logs lower than HMV isolates (Figs.3D, 3E and 3F), indicating that HMV *K. pneumoniae* strains were more virulent than non-HMV isolates.

**FIG 3.**
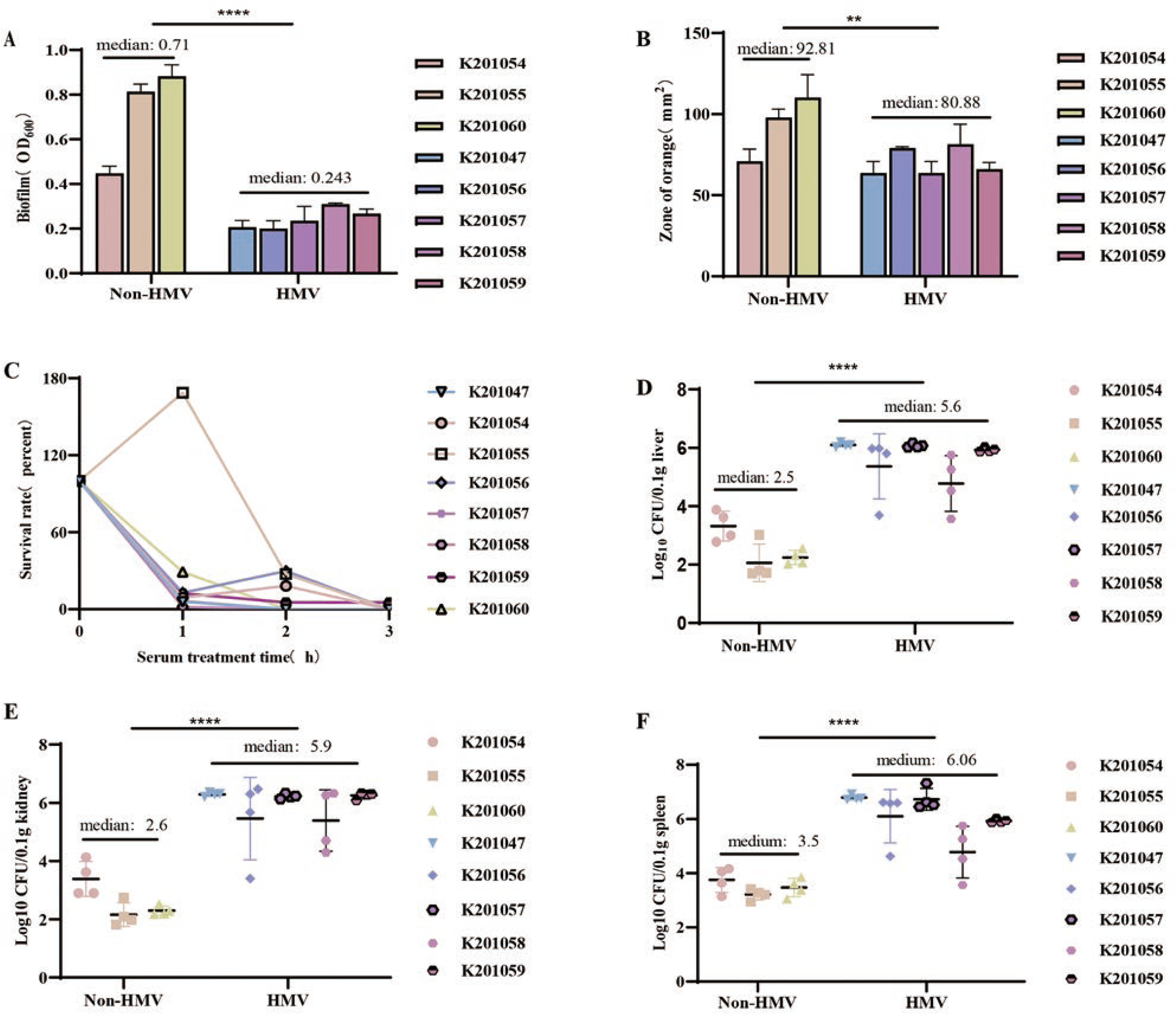
The microbiological phenotype of eight isolates. (A) Analysis of biofilm formation. **** *p* < 0.0001 by *t* test. (B) Siderophore production assessed by the CAS assay. ** *p* < 0.01 by *t* test. (C) Serum killing assay. (D) Colonization levels in the liver of infected mice. (E) Colonization levels in the spleen of infected mice. (F) Colonization levels in the liver of infected mice. Bacterial numbers (expressed as log_10_ CFU) were standardized per 0.1g of wet organ weight. The unpaired nonparametric Mann-Whitney U test was used to test for statistical significance of the infection test in mice. **** *p* < 0.0001.

### WGS revealed that the eight isolates belonged to the same clone

To investigate the genomic characteristics of the co-occurrent HMV and non-HMV *K. pneumoniae* isolates, we performed whole genome sequencing (WGS) on the eight strains recovered. Analysis of genomic data revealed that all strains belonged to the K57-ST412 lineage, with an average genome size of 5.6 Mb, a GC content of approximately 57-58%, and average nucleotide identity (ANI) values ranging from 99.96% to 99.99% between isolates. These results confirmed the intimate genetic relatedness of the eight isolates. Pulsed-field gel electrophoresis (PFGE) profiling divided the eight isolates into two main subtypes, one composed by the non-HMV isolates and strain K201059, and the second formed by the remaining HMV strains (Fig.S1).

Analysis of plasmid content with PlasmidFinder revealed the existence of two putative plasmids, which were designated as pA and pB, respectively. pA contained an IncFIB(k) replicon, and exhibited a high similarity with pGN-2, a plasmid harbored by the *K*. *pneumoniae* strain GN-2(99.86% identity and 68% coverage, accession no. NZ_CP019161.1), albeit with a higher number of transposases and hypothetical protein coding genes. The pB plasmid contained a Col440II replicon with high homology to pBio19 found in Turkey and Iraq (99.98% identity and 100% coverage, accession no. NZ_CP096811.1) (Fig.S2). BLAST analysis against the virulence genes database revealed the isolates contained several virulence determinants, including the *rmpA* gene that regulates the mucoid phenotype, the *wabG* gene that participates in the biosynthesis of the core lipopolysaccharide, genes for the biosynthesis or uptake of iron (*iutA, iroBCDN*, *irp2* and *ybts*) and fimbrial genes (*fimA, marKA* and *markD*).

### Isolates contain mutations in metabolic and regulatory genes

To gain an insight on the genomic determinants behind HMV development in co-occurring *K. pneumoniae* isolates, we used the genome structure of K201047 as a reference and performed a whole-genome alignment of all other seven isolates against the K201047 genome. Our analysis identified 193 mutations events (SNPs and InDels), most of which were either located in intergenic regions (119 mutations) or were synonymous mutations (62 mutations), with only 12 mutations being non-synonymous mutations. And these non-synonymous mutations preferentially occurred in genes encoding regulators and metabolism (Table 1). Subsequently, to explore the genetic determinants associated with the HMV phenotypic changes, we first focused on the non-synonymous mutations that occurred in the non-HMV strains and not in the HMV strains. Firstly, we identified a T-base deletion that disrupts the *rmpA* gene, a regulator of mucoid phenotype, which predicted to be responsible for the non-HMV phenotype in strain K201054. In the other two non-HMV isolates, K201055 and K201060, they both carry a single missense mutation in the *wbaP* gene(S435R), a undecaprenyl-phosphate galactose phosphortransferase which involved in the complex *cps* synthesis. To confirm that mutations in *rmpA* and *wbap* were involved in the development of non-HMV phenotype, we constructed complemented plasmids containing the CDS region and its native promoter. Introduction of the native *rmpA* gene into the K201054 non-HMV strain restored the HMV phenotype (Fig.S3). On the contrary, introduction of the native *wbaP* gene into non-HMV strains K201055 and K2010560 was unsuccessful. Yet, the deletion of the *wbaP* gene has previously been confirmed to impair the production of capsule, corresponding with a non-HMV appearance on plate(12, 13). Thus, mutation in *wbap* appears to be a mechanism employed by strains K201055 and K201060 that induced the formation of the non-HMV phenotype. Besides, we also observed multiple independent mutations in the genes *recC*, *minE*, *ydcJ* and *hrpB*, the latter encoding an ATP-dependent helicase that is known to promote pathogenicity as an effector molecular(14). To the best of our knowledge, these four genes have not been documented to be associated with the HMV phenotype, and additional research is needed to investigate their potential contribution.

**Table 1.**
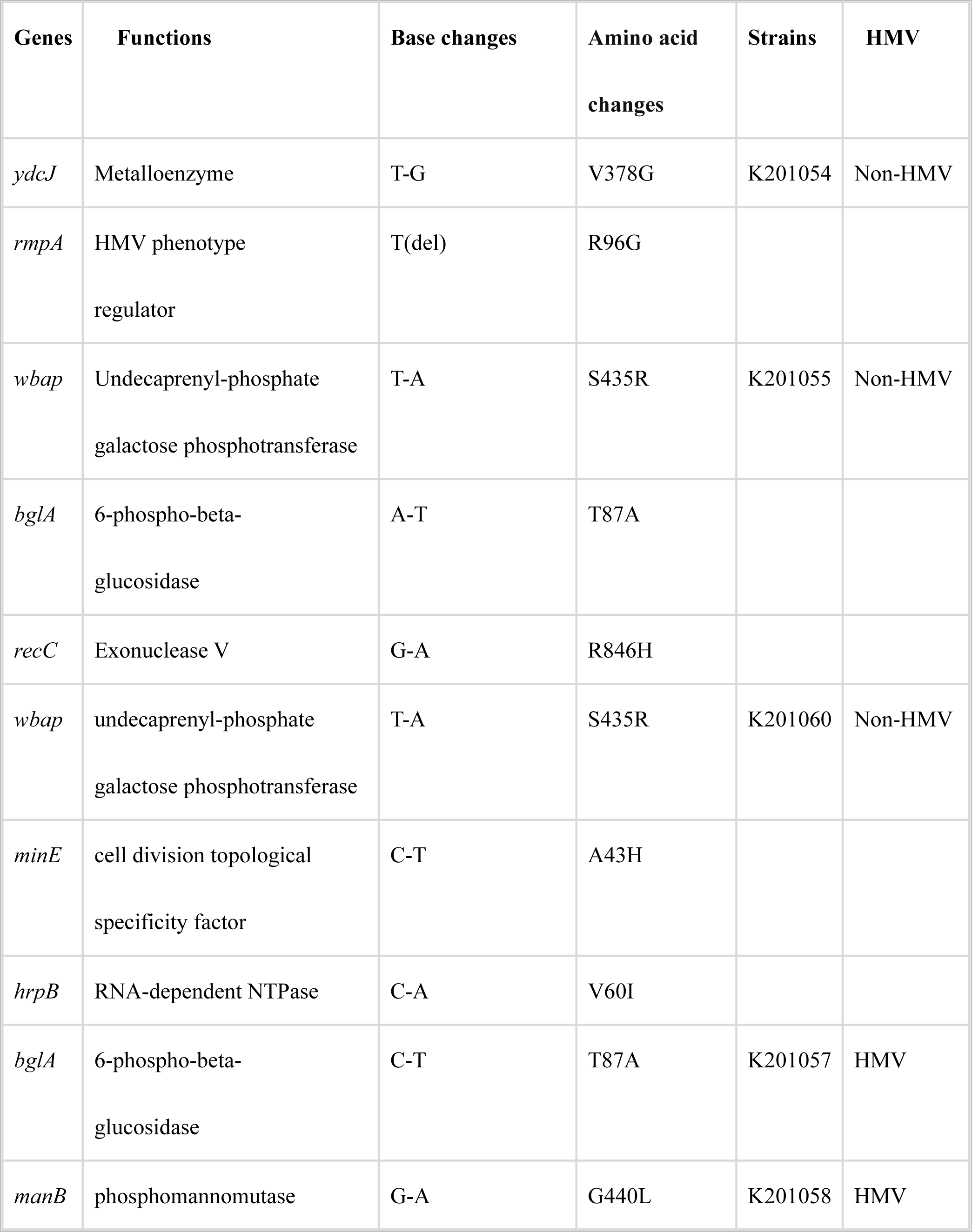
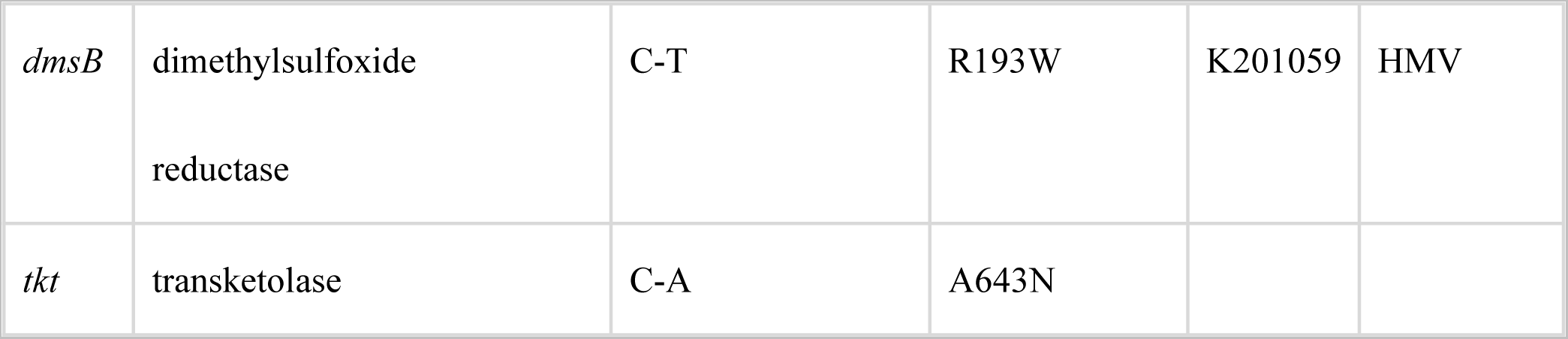
Non-synonymous mutation and InDels found in the strains

For the clinical HMV isolates K201057 and K201059, firstly, two distinct mutations relating to carbohydrate metabolic pathways—the *tkt* gene linked to glycolysis and *bglA* implicated in the phosphorylated disaccharide metabolism pathway—were found. Additionally, we also observed one independent mutation in the gene *dmsB* in the HMV isolate K201059, encoding the electron-transfer subunit of the dimethylsulfoxide reductase(15). Subsequently, we noticed a base substitution in *manB* distributed at *cps* locus in the HMV strain K201058, which is implicated in convergent signal transduction of capsule biosynthesis. It is well known that HMV and the *cps* locus involved in capsular production are functionally associated (16). Here, the mutation of *manB* did not affect the HMV phenotype of strain K201058.

### Differentially expressed genes involved in transport and metabolic activities are associated with non-HMV phenotypes

Isolates K201059(HMV) and K201060(non-HMV) were selected for a genome-wide RNA-Seq analysis to identify differences in their transcriptomic profiles potentially contributing to their distinct phenotypic characteristics. A total of 450 differentially expressed genes(DEGs) were identified in strain non-HMV K201060 compared to HMV strain K201059(|log2 (fold change) |>1, *p*<0.05), with 71 up-regulated transcripts (15.7%) and 379 down-regulated transcripts (84.3%). As shown in Fig. 4A, the most significantly up-regulated gene was found to be *cysK*, encoding the cysteine synthase A, and the most significantly down-regulated gene was *vgrG*, encoding the homonymous type-VI secretion protein, with log_2_FC 3.4 and -7.94, respectively. Subsequently, we selected 13 DEGs to validate the RNA-Seq data via RT-qPCR analyses, which highly correlated with the RNA-Seq data (Fig. 4B).

**FIG 4.**
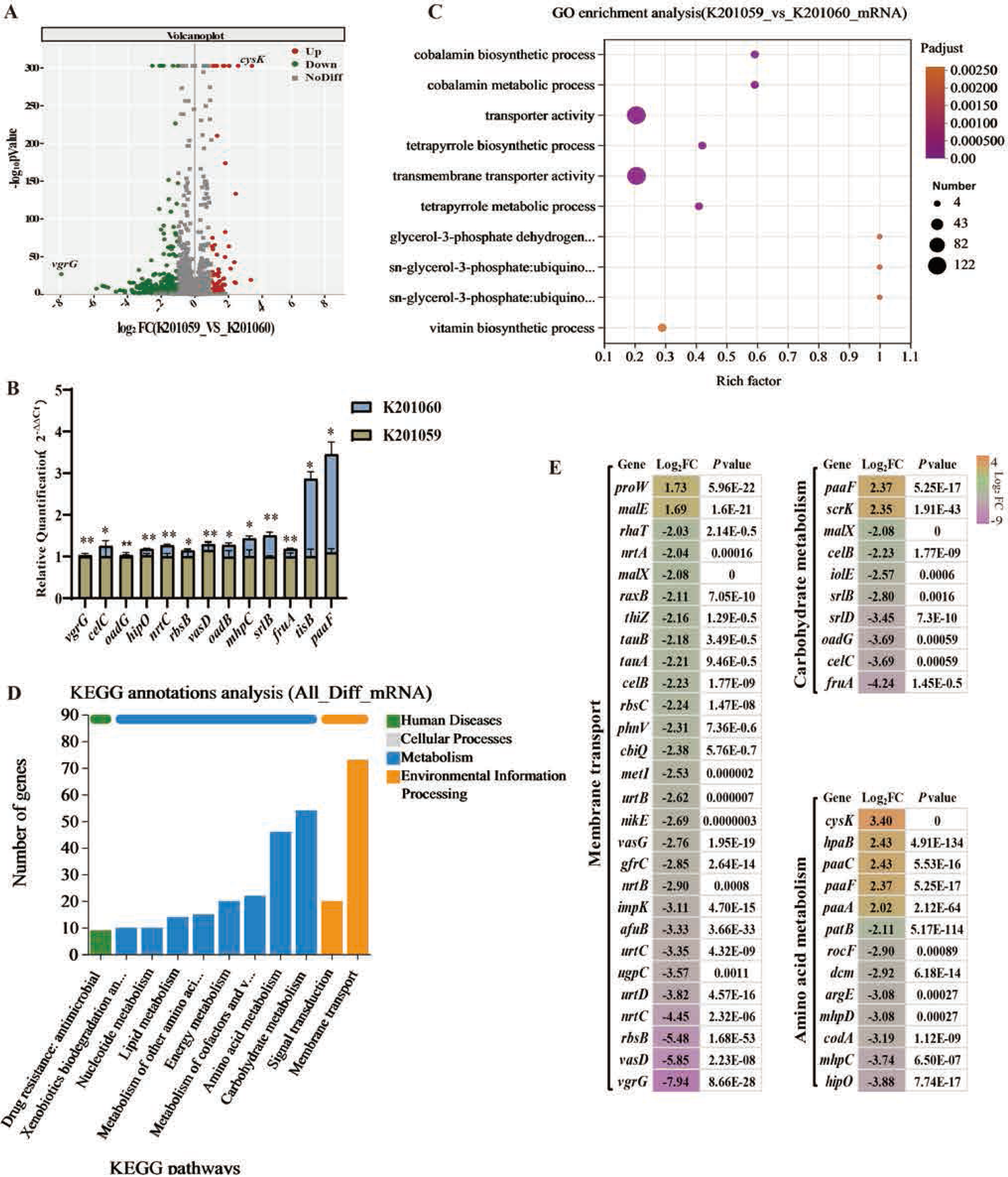
RNA-seq gene expression profile of non-HMV isolate K201060 compared with HMV isolate K201059. (A) The volcano plot of DEGs in bacterial cells. -log10pvalvue indicates the statistical test value for the difference in gene expression change, that is, the p value. (B) RNA-Seq validation by RT-qPCR experiment. (C) GO enrichment analysis of DEGs. Rich factor refers to the number of genes annotated to the GO term in a gene set. (D) Clusters of KEGG pathways. (E)Map of typical DEGs associated with transmembrane transport activity, carbohydrate metabolism and amino acid metabolism.

A gene enrichment analysis was conducted to identify dysregulated gene functions and biological pathways. The differentially expressed genes were mapped to Gene Ontology(GO) categories and KEGG pathways. We found that the DEGs were clustered within 10 GO functional categories, with “transporter activity” and “transmembrane transporter activity” dominating the GO categories(Fig.4C). Subsequently, a KEGG enrichment analysis was further performed to categorize DEGs into diverse pathways. As shown in Fig.4D, the results indicated that membrane transport, carbohydrate metabolism and amino acid metabolism were the most enriched pathways in DEGs, and other major pathways included metabolism of cofactors and vitamins, energy metabolism and cellular community-prokaryotes processes.

Based on GO and KEGG analysis, it was found that 26 of the 28 DEGs associated with membrane transport activity were downregulated. In particular, *vgrG* and *vasD* involved in the type VI secretion system, *rbsB* encoding the ribose ABC transporter and *nrtC* encoding the nitrate ABC transporter were significantly down-regulated in non-HMV isolate K201060 (|log2 (fold change) |>4). Besides, the expression of eight of the ten genes related to carbohydrate metabolism were also down-regulated including the PTS fructose transporter(*fruA*), sorbitol transporter (*srlB*), PTS family sugar-specific enzyme(*celC*) and oxaloacetate decarboxylase(*oadG*). Interestingly, RNA-Seq data showed that genes participating in cysteine metabolism (*dcm*, *cysk*, and *patB*), genes participating in phenylalanine metabolism (*hipO*, *paaA/C/F*, and *mhpC/D*), and genes participating in arginine metabolism (*argE*, *codA*, and *rocF*), were also significantly dysregulated(Fig.4E).

## Discussion

As a preponderant bacterium driving community-acquired infections, *K. pneumoniae* with HMV has the ability to cause complicated infections, including pneumonia, liver abscesses, and sepsis(17). Clinical *K*. *pneumoniae* can undergo HMV phenotypic variation in response to diverse hostile environments, with the result that both HMV and non-HMV phenotypes can be observed within the same clonal population(7). Published data for *K*. *pneumoniae* ST11 strains demonstrated that switching of the HMV phenotype can provide a fitness advantage for invasive infections(18). In this study, we observed that HMV and non-HMV phenotypes co-occurred in the same clonal isolates. We investigated the underlying mechanisms using whole genome sequencing and RNA-Seq profiling. We speculate that mutation in *rmpA(*R96G) was responsible for the non-HMV phenotype of strain K201054, in agreement with evidence associating *rmpA* and the HMV phenotype of *K*. *pneumoniae* has been confirmed(19, 20). Furthermore, the mutation S435R in *wbap* reduced the overproduction of capsule polysaccharide in the non-HMV strains K201055 and K201060. A strong link between capsule production and the HMV phenotype has been reported, despite a few reports suggesting that HMV does not require overproduction of capsule(16). We confirmed a reduction in CPS synthesis in clinical isolates K201055 and K201060, corresponding to a non-HMV appearance on the plates.

Membrane transporters play a key role in play a key role in controlling physiological functions such as mediating the flow of molecules between cytosol and the extracellular environment, including energy sources(21). In the present study, the expression of most genes participating in transporter activity was down-regulated in a model non-HMV isolate, suggesting that membrane transporter activity may be another component of the complex CPS biosynthesis and HMV regulatory networks and possibly coordinating these features with the intracellular transport of substances or energy metabolism. In addition, we confirmed that the expression of genes involved in carbohydrate metabolism (*fruA*, *celC* and *oadG*) and amino acid metabolism (*hipO*, *mhpC* and *codA*) was also down-regulated. Carbohydrate metabolism and amino acid metabolism are essential for nutrient acquisition(22, 23). CPS biosynthesis and HMV are in fact extraordinarily energy-intensive processes, so it is intuitive that disruption or repression of genes involved in carbon metabolism may result in reduced CPS biosynthesis and HMV. This supports the notion that the cellular metabolic status in relation to nutrient availability could be a conserved mechanism in CPS synthesis and HMV.

Mutations are not necessarily random, but may be the result of a strong genotype-by-environment interaction that enhance recombination function, which is also a manifestation of adaptation effects under appropriate selective pressure(24). In the clinical isolates retrieved in our study, a total of 192 mutations were identified, of which the majority of the identified mutations events (SNPs and InDels) identified were located in the intergenic regions or were synonymous mutations, with only 12 mutations being non-synonymous mutations. Here, our mutation studies focused on the coding regions, this may result in the association of some intergenic regions of mutations with the HMV phenotype being excluded. Nevertheless, intergenic mutations may actually affect the transcriptional activity of genes involved in host interaction, metabolism and antibiotic susceptibility, identification of the mutations in intergenic regions is also important(25). Further studies, particularly of upstream region of genes with unknown functions, will likely uncover novel mechanisms of HMV phenotypic switching.

In summary, our work elucidates that the co-occurrence of HMV and non-HMV phenotypes in the same clonal isolate may be driven by mutational mechanisms, as well as down-regulation of genes involved in metabolism and transporters, with a substantial impact on the ability to form biofilm.

## Materials and Methods

### Antimicrobial susceptibility testing

All isolates were examined for antimicrobial susceptibility testing (AST) using broth microdilution following the standard Clinical and Laboratory Standards Institute (CLSI) guidelines. Isolates were tested for ceftazidime, cefotaxime sodium, ciprofloxacin, streptomycin, chloramphenicol, tetracycline, imipenem, meropenem, gentamicin, polymyxin. E. coli ATCC 25922 and *K. pneumoniae* ATCC 700603 were used as the quality control strains, and the interpretation of the results was based on 2020 CLSI M100 30th Edition break-points, which referred to the EUCAST v.11.0 breakpoints (https://eucast.org/).

### Biofilm assays

Biofilm formation assays were performed as described elsewhere(26). Briefly, 200 µL of bacterial cture (1.5 × 10^7^ CFU/mL) was added to wells in 96-well flat-bottomed polystyrene plates, and incubated for 18 to 24 h at 37°C. At the end of incubation period, planktonic bacteria were removed by washing thrice with 200 µL of distilled water. Wells were dried, treated with 200 µL of 0.5% crystal violet stain for 20 minutes, and washed thrice with distilled water. The bound dye was solubilized with 200 µL of 36% glacial acetic acid and quantified by measuring the optical density at 600 nm (OD_600_).

### Serum resistance assay

Bacterial suspensions containing 1.5 × 10^6^ CFU/mL were collected from mid-log phase cultures, mixed at a 1:3 (vol/vol) ratios with nonimmune human serum donated by healthy volunteers, and incubated at 37°C. The colony count is determined by serial dilution method at 1,2 and 3h(27). Resistance grading was defined from grade 1 to grade 6.

### CAS agar assays for iron uptake

To elucidate the ability of HMV isolates and HMV isolates to secrete siderophere, we prepared CAS agar plates as described in a previous publication. Then, isolates were prepared with a concentration of 1.5 × 10^8^ CFU/mL, 5µL of bacterial cultures were inoculated on CAS plate and cultivated at 37℃ for 48h. The orange halo produced on the CAS plates was measured. The assay was repeated three time(28, 29).

### String test, mucoviscosity testing and quantification of polysaccharides

The HMV phenotype relies on the classical string test as described (30, 31). The mucoviscosity levels were determined by natural sedimentation assay. Briefly, isolates were cultivated in M9 at 37 °C overnight. The following morning 1 mL of optical density OD_600_ normalized bacteria was precipitated by the action of gravity for 24h. The absorbance of the supernatant was measured at OD_600_. Capsule polysaccharide(CPS) was extracted according to previously described methods(32). The quantification of CPS was determined from a standard curve of D-glucuronic acid and expressed as micrograms per 10^6^ CFU.

### PFGE profile

The assay was performed as described previously(33). DNA was digested with XbaI (60U) for 4h before the digested DNA was separated in a 1% agarose gel (6V/cm, 17.5h). Bionumerics software (UPGMA clustering method) was used for isolate profiling.

### Mice infection experiments

We used an intraperitoneal infection model to investigate the pathogenicity of the distinct bacterial isolates. Five-week-old female BALB/c mice were intraperitoneally injected with 5.5 X 10^7^ CFUs in 200ul of phosphate buffer solution (PBS). The mice were euthanized at 24 h postinoculation (hpi) and the amounts of bacteria in liver, spleen, and kidneys were determined.

### Whole Genome Sequencing(WGS) and Single Nucleotide Polymorphism(SNP) Analysis

All isolates were sequenced using an Illumina HiSeq to generate 150 bp pair-edend reads and 100x coverage, while K201054 strain was analyzed using PacBio RSII analysis to obtain complete chromosome and plasmid sequences (Majorbio Co., Ltd. Shanghai, China). We used SOAPdenovo v.2. and Unicycler v0.4.6 for *de novo* assembly and obtaining of the complete genome sequence(34). Gene annotations were carried out with the RAST online tool(35) (http://rast.theseed.org/FIG/rast.cgi). The MLST profile was assigned according to the online database of MLST v.2.0, while the capsular type of *Klebsiella* strains was determined according to wzi gene sequence. Plasmid replicon types were identified by the server Plamid Finder(https://cge.foof.dut.dk/services/PlasmidFinder)(36). Similarity of WGS were assessed by the JSpeciesWS Online Server(37) (https://jspecies.ribohost.com/jspeciesws). The circular map generated using CGView(38)(https://paulstothard.github.io/cgview). Virulence determinants were determined through the VFDB server(39) (http://www.mgc.ac.cn/VFs/main.htm). Single-nucleotide polymorphism (SNP) analysis was performed by the kSNP 3.0 based on concatenated genome sequence data of eight *K. pneumoniae* ST412 isolates(40). The mutation sites were annotated by snpEff (https://snpeff.sourceforge.net/SnpEff.html).

### Construction of recombinant strains

Construction of recombinant plasmids for gene complementation was performed as described previously(41). The target gene was amplified by PCR and inserted into the shuttle vector pBBR1MCS-3(see Table S1 in Supplemental materials for primer). After that, the recombinant plasmid was transformed into the target isolates by conjugation.

### RNA sequencing and analysis

HMV strain K201059 and non-HMV strain K201060 were grown overnight in LB medium. The bacterial overnight cultures were diluted at 1:100 in fresh LB medium and grown to a mid-exponential phase at 37°C. The bacteria were harvested by centrifugation and total RNAs were extracted using a Spin Column Bacteria Total RNA Purification kit (Sangon Biotech) according to the manufacturer’s instructions. The RNA-Seq were performed by the Majorbio Bio-pharm Technology Co. Ltd (Shanghai, China) with the Illumina Hi Seq X Ten platform. Gene expression difference analysis was performed by DESeq2. Genes with p-adjust < 0.05 or log_2_FC >1 were considered to be significantly differentially expressed genes(DEGs). GO enrichment analysis was performed with top GO, and KEGG pathway and COG classification enrichment analysis were performed by cluster profile, respectively.

### Quantitative reverse-transcription PCR(RT-qPCR)

The RT-qPCR was performed as described elsewhere to verify the mRNA levels of differentially expressed genes in the transcriptome(42). The isolates were grown overnight in LB medium. The bacterial overnight cultures were diluted at 1:100 in fresh LB medium and grown to mid-exponential phase at 37 ℃ . Ttotal RNA was harvested from the isolates using the Spin Column Bacteria Total RNA Purification KIT (Sangon Biotech, Shanghai). Then, the extracted RNA was reverse transcribed into cDNA using a TransScript All-in-One First-Strand cDNA Synthesis SuperMix (One-Step gDNA Removal). Finally, the RT-qPCR was performed using a Tip Green qPCR SuperMix (TransGen Biotech Co., Ltd.) in a Mastercycler ep realplex system (Eppendorf, Hamburg, Germany), with an initial incubation at 94°C for 30 s, followed by 40 cycles of 5 s at 94°C and 30 s at 60°C. The internal control gene 16SrRNA was used to normalize the expression of each transcript (see Table S1 in Supplemental materials for primer).

### Statistical analyses

Statistical analyses were performed with GraphPad Prism software Version 8. *K*. *pneumoniae* burden in tissues were examined based on unpaired nonparametric Mann–Whitney U tests. Other phenotypic assays were analyzed by *t* test. Statistically significant was defined by P<0.05 (*), P<0.01 (**), P<0.001 (***), and P<0.0001 (****).

### Ethics statement

This study has obtained ethical approval and imformed consent of the patient. The ethical code is 2022Y – 194. Furthermore, the animal experimental protocols were approved by the Ethics Committee of Experimental Animals, Southwest Medical University. The approval number was 20210225-2.

### Data availability

The complete sequences of the isolate in this study, K201054, were deposited in the GenBank databases (accession number. CP110637-CP110639). The draft genome sequences of other seven isolates were submitted to Genbank under accession numbers JAPJTY000000000, JAPJTX000000000, JAPJTZ000000000, JAPJUA000000000, JAPJUB000000000, JAPJUC000000000 and JAPJUD000000000, respectively. RAW sequence reads are available on NCBI under Bioprojiect accession number PRJNA891137.

## ACKNOWLEDGMENTS

This research was supported by the Sichuan Province Science and Technology project [2022YFS0631 and 2022YFS0632], the Natural Science Foundation of Luzhou (2021-NYF-20), the Joint Project of Southwest Medical University [2021SNXNYD01 and 2021NJXNYD06], and the Southwest Medical University Foundation (2021ZKZD002).

## Supplementary Material

### Supplementary Figures

**Figure S1.** PFGE cluster analysis showed that the 8 isolates could be classified into 2 subtypes (A1 and A2).

**Figure S2.** Circular map of the plasmid pA and plasmid pB and comparative genomics analysis with its similar plasmids. Gene annotation. Red, virulence-related genes.

**Figure S3.** String test for isolate K201054. (A)The string test revealed that K201054, harboring a mutation in rmpA, was non-HMV by string test. (B) Complementation of the native *rmpA* gene into the K201054 non-HMV strain restored the HMV phenotype (K201054-p*rmp* strain).

### Supplementary Table

**TABLE S1.**
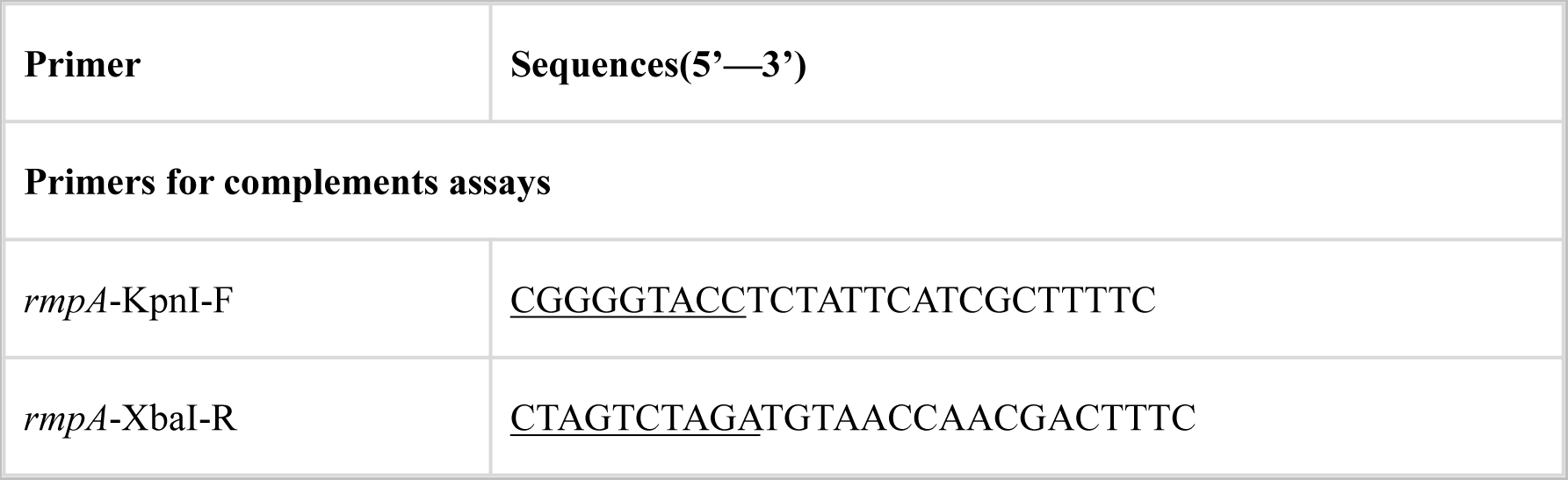
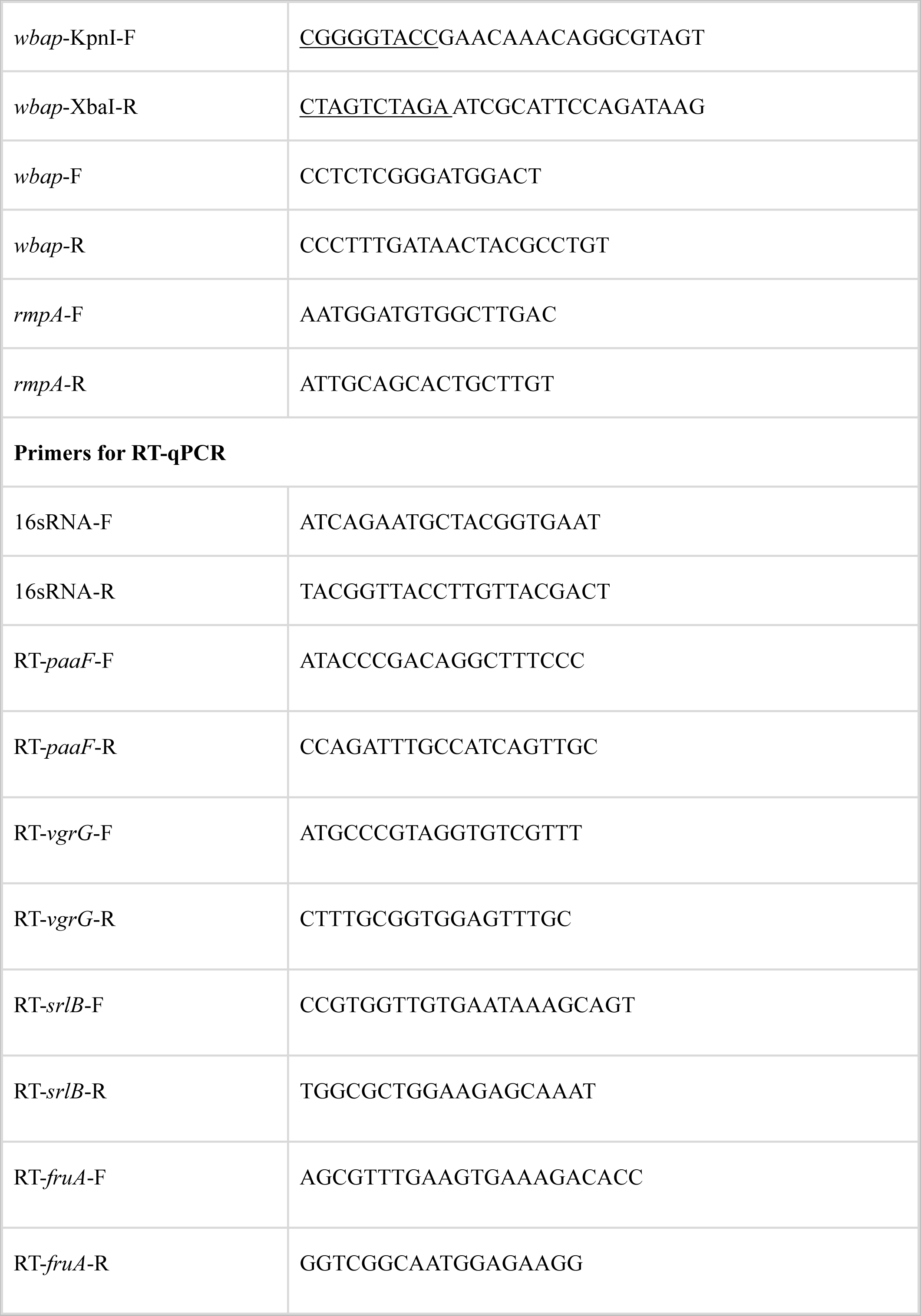
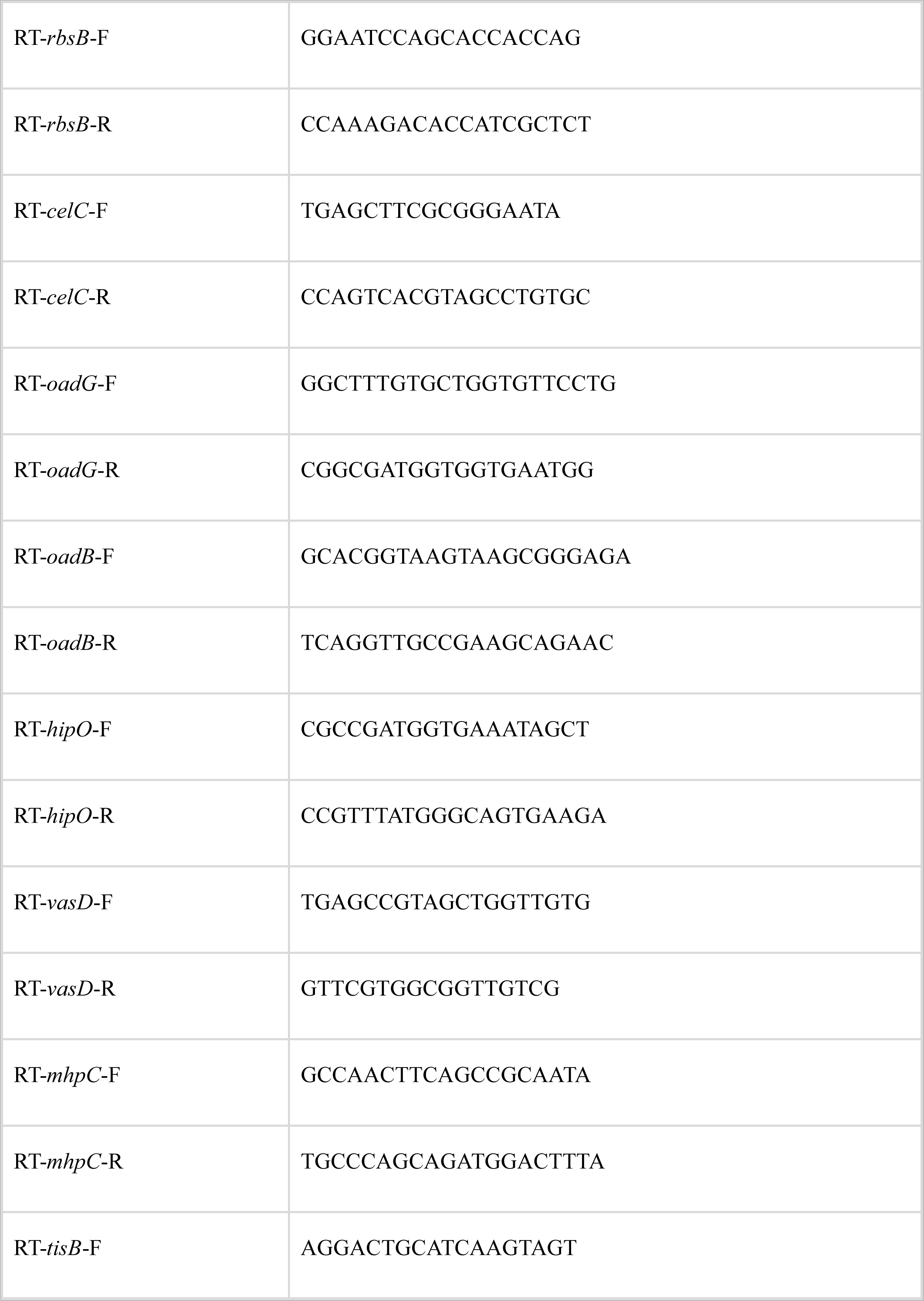
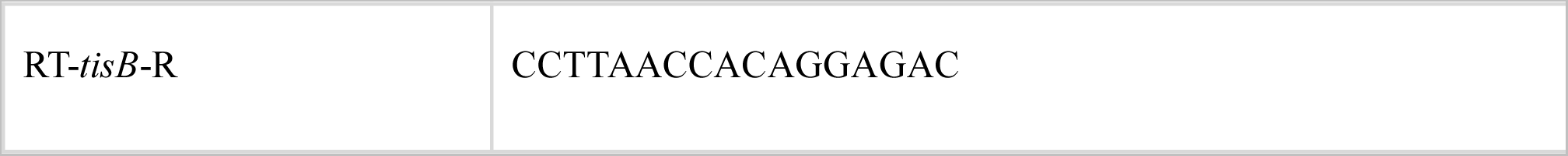
Primers used in this study

